# Effects of land use change and elevation on endemic shrub frogs in a biodiversity hotspot

**DOI:** 10.1101/2024.05.17.594636

**Authors:** Himanshu Lad, Ninad Gosavi, Vijayan Jithin, Rohit Naniwadekar

## Abstract

Agroforestry, often promoted as a sustainable agriculture practice, is rapidly expanding, often at the cost of primary tropical forests. While agroforestry impacts amphibian diversity negatively, its effects on population demography, microhabitat use and body condition are relatively understudied. This information is crucial for determining and promoting amphibian-friendly land use practices. We compared habitats, population densities, microhabitat use, and body condition of two endemic species of shrub frogs (*Pseudophilautus amboli* and *Raorchestes bombayensis*) across 1) elevations (low- and high-elevation forests) and 2) land use categories (cashew, rubber, and low-elevation forests) in the northern part of the Western Ghats Biodiversity Hotspot. Using distance sampling, we demonstrated that abundances of the two shrub frog species differed across elevation categories, with *Pseudophilautus* more common in low-elevation forests and *Raorchestes* more prevalent in high-elevation forests. Both species of frogs exhibited extremely skewed, male-biased sex ratios, with three females for 100 males. *Pseudophilautus* had lower densities and poor recruitment and exhibited altered microhabitat use in cashew plantations compared to low-elevation forests. Although adult male *Pseudophilautus* densities in rubber were similar to those in low-elevation forests, they exhibited altered microhabitat use and smaller body sizes than in forests, indicating poor body condition. We demonstrate differential impacts of agroforestry types on shrub frogs. We also demonstrate that distance sampling can be a useful tool for population monitoring of shrub frogs, which comprise almost 25% of the anuran diversity in the Western Ghats. Additionally, there is a need to identify the drivers of extremely skewed sex ratios, which make these species vulnerable to population crashes. Considering the recent downlisting of the two focal species to Least Concern, we advocate for their uplisting to at least Near Threatened status in light of their patchy distribution, negative impacts of rapidly expanding agroforestry plantations and extremely skewed sex ratios.

## INTRODUCTION

With the human population expected to reach 9.5 billion by 2050, food production will need to increase by 60% to meet growing demands. In response, agroforestry is often advocated as a sustainable alternative for better land management and production improvement (FAO, 2022). Agroforestry is also positioned as a vital climate solution, as it is believed to aid in restoring degraded lands, halting forest degradation, and contributing to biodiversity conservation (FAO, 2024). However, in many tropical regions, forests are being converted into agroforestry plantations (Rege & Lee, 2022), leading to forest and biodiversity loss. Furthermore, the impacts of agroforestry plantations on biodiversity vary. While the response of bird and reptile richness depends on the type of agroforestry, amphibians consistently experience negative impacts (Palacios, Agüero, & Simonetti, 2013; Bohada-Murillo, Castaño-Villa, & Fontúrbel, 2020; Cervantes-López & Morante-Filho, 2024). With 40.7% of species being threatened, amphibians are the most imperilled vertebrate group, marked by a rising trend of documented extinctions (Luedtke et al., 2023). Among frogs with different modes of development, direct-developing frogs are particularly vulnerable (Murray, Nowakowski, & Frishkoff, 2021). Agriculture ranks among the most pervasive drivers of amphibian loss (Fulgence et al., 2022; Luedtke et al., 2023). Yet, relatively fewer studies have focused on assessing species responses to different forms of land use, particularly concerning amphibians (Palacios, Agüero, & Simonetti, 2013). While some recent studies have examined the effects of agroforestry plantations on species richness and composition (Trimble & van Aarde, 2014; Pereyra et al., 2018; Fulgence et al., 2022), fewer have investigated its impacts on population demography, microhabitat use, and body condition (Bodinof Jachowski & Hopkins, 2018). Furthermore, while considerable information exists on amphibian responses to habitat conversion involving cacao, coffee, oil palm, pine, and eucalyptus, information from other forms of agroforestry, like cashew and rubber, remains limited (Palacios, Agüero, & Simonetti, 2013). This information is critical for guiding agroforestry plantation management to promote biodiversity conservation. Given this context, it is imperative to evaluate the impacts of agroforestry on direct-developing amphibians.

Population monitoring of amphibians in light of threats posed by land use and climate change and emerging diseases has been identified as a crucial measure for periodic amphibian conservation assessments and priortisation (Luedtke et al., 2023; Re:wild, Synchronicity Earth, 2023). Methods that allow reliably estimating and monitoring populations for a diverse array of species with modest efforts can be valuable. Monitoring amphibians across gradients typically focus on comparing capture or encounter rates. However, these measures may not fully account for variation in detection probability, which is particularly relevant for small-sized, cryptically coloured arboreal frogs concealed within vegetation in natural environments (Mazerolle et al., 2007). Alterations in undergrowth in modified habitats can result in varying detection probabilities across land use types, underscoring the importance of employing appropriate methods such as mark-recapture or distance sampling to address this variability in detection probability. Distance sampling has proven effective in estimating frog densities across habitats (Pikacha et al., 2016). Unlike mark-recapture, distance sampling is less time-intensive and does not involve handling frogs (Fogarty & Vilella, 2001; Funk et al., 2003). Additionally, it is a valuable tool for reliably estimating animal population densities, aiding in long-term monitoring of amphibians and periodic conservation assessments, which is critical for guiding conservation action. Therefore, there is a need for determining the utility of the method for monitoring frogs.

The Western Ghats of India is a crucial part of the Western Ghats - Sri Lanka Biodiversity Hotspot, which is recognised as one of the hottest biodiversity hotspots (Myers et al., 2000). Despite its importance, less than 10% of the Western Ghats is within the Protected Area network, leaving areas outside vulnerable to high rates of forest loss and degradation (Reddy, Jha, & Dadhwal, 2016). This trend is particularly pronounced in the northern Western Ghats, where Protected Areas are primarily confined to higher elevations, leading to extensive habitat conversion to agroforestry plantations (Kulkarni & Mehta, 2013; Rege, Warnekar, & Lee, 2022). Such conversions have detrimental impacts on biodiversity, especially at lower elevations (Jithin, Rane, Watve, Giri, et al., 2023; Biswas et al., 2023, 2024), underscoring the necessity of comparing species responses to different land use types. The Western Ghats harbour an exceptional diversity of amphibians, with over 232 species of frogs, 69% of which are endemic to the region and threatened (Luedtke et al., 2023; Re:wild, Synchronicity Earth, 2023). Among the different anuran clades, shrub frogs exhibit remarkable diversity and endemism, with more than 60 species documented across the Western Ghats (>25% of known species of frogs). Many of these shrub frogs have narrow geographic ranges and are threatened by habitat loss and climate change (Vijayakumar et al., 2016; Sankararaman & Miller, 2024). These frogs employ direct development as a mode of reproduction, laying their eggs in leaf litter or on moss-laden tree trunks, where they hatch directly into small froglets. Focusing on two such species, *Pseudophilautus amboli* and *Raorchestes bombayensis*, our study aims to shed light on their population demography and habitat utilisation across different land use types. *Pseudophilautus amboli* (hereafter *Pseudophilautus*) is endemic to the northern and central Western Ghats of India, and *Raorchestes bombayensis* (hereafter *Raorchestes*) is restricted to the northern Western Ghats. Although recently downlisted from Critically Endangered and Vulnerable to Least Concern, this decision was made with limited ecological information on these species (IUCN SSC Amphibian Specialist Group, 2020, 2021). The lack of data on population demography and species responses to land use change underscores the importance of our research in informing conservation efforts for these shrub frogs.

To this end, we asked the following questions: 1) How do microhabitats for shrub frogs vary across different land use categories? 2) What are the population densities of adult and juvenile shrub frogs in the different land uses? 3) How do shrub frogs utilise micro-habitats across different land uses? and 4) Is there variation in body size of adult male shrub frogs across different land use categories? Considering that sympatric shrub frogs may differ in preferred environmental conditions, we expected differences in population densities of the two species across elevation categories (hypothesis 1). We further expected that human modification in agroforestry plantations (rubber and cashew) would impact microhabitat availability for shrub frogs (hypothesis 2). Given the alterations in habitat and the cascading effects, we expected differences in population densities of adult and juvenile shrub frogs (hypothesis 3) and their microhabitat use (hypothesis 4) across differ across land use types. Given the changes in microhabitat availability and use, we expected alterations in the body condition of shrub frogs (hypothesis 5).

## SAMPLING METHODS

### Study Area

We conducted the study from June to September 2022 in the northern portion of the Western Ghats of India coinciding with peak monsoon in the region (June to October). Given the high endemism and threats posed by anthropogenic activities, the Western Ghats is one of the “hottest” hotspots (Myers et al., 2000). Over 85% of the 450 amphibians recorded in the Western Ghats are endemic to the region (Dahanukar & Molur, 2020). More than 67 species of arboreal frogs (Family: Rhacophoridae) occur in the Western Ghats, most of which are endemic to the region (Dahanukar & Molur, 2020). The diversity of arboreal frogs is lower in the northern Western Ghats due to environmental and historical factors (Daniels, 1992; Cyriac et al., 2024).

We conducted our sampling in the Dodamarg and Sawantwadi Taluks of the Sindhudurg District in Maharashtra (15°40’–16°0’N; 73°51’–74°9’E). The region experiences an annual rainfall ranging from 2,300 to 3,200 mm, with the peak rainfall typically observed between June and September during the south-west monsoon season. The elevation across our sampling sites ranged from 50 to 900 m above sea level.

Most of the tropical forests in the lower elevations are privately owned, except for a few patches of Reserved Forests under the jurisdiction of the Sawantwadi Forest Division. Under Indian forest laws, local communities have access to Reserved Forests to non-timber forest produce collection and cattle grazing. The Forest Department may conduct timber felling in these forests according to their working plans (Anonymous, 1972). Vegetation in the lower areas is typically classified as moist-deciduous and semi-evergreen, and the higher elevations harbour evergreen forests (Jog, 2009; Tadwalkar et al., 2020). However, a recent study suggests chronic human disturbance has converted evergreen forests into deciduous forests across low and high elevations (Biswas et al., 2024). The area has witnessed significant conversion of private forests into cashew, rubber, and pineapple plantations in recent decades (Rege & Lee, 2022; Munje & Kumar, 2022; Biswas et al., 2023). Within our study area, five species of Rhacophorid frogs are found, including the three species of shrub frogs - *Pseudophilautus amboli*, *P. wynaadensis*, and *Raorchestes bombayensis* - although *P*. *wynaadensis* is extremely rare (Katwate & Apte, 2019; Komanduri, Sreedharan, & Vasudevan, 2023).

### Shrub frog density

We marked 28 transects across the four categories: low- and high-elevation forests, cashew and rubber (Table S1). The minimum and maximum length of transects walked varied between 230 and 300 m, respectively (Table S1). We monitored most transects four times during the study duration (June and September 2022). However, we could monitor two transects only three times due to the presence of wild elephants. Two observers (HL and NG) walked the transects between 1830 and 2330 hr, coinciding with peak shrub frog activity. We scanned for frogs using a headlamp and handheld torch. Upon spotting a frog, we recorded the species identity, number of individuals, and the perpendicular distance of the shrub frog sighting, following the conventional distance sampling protocol (Buckland et al., 2005). During the monsoon, shrub frogs perch on leaves, branches, and tree trunks (Fig. 1). We classified the animals as adult males, adult females and juveniles. Adult males were identified based on their vocal sacs, females were identified based on their larger size and/or absence of vocal sacs and/or presence of eggs in the body, and juveniles were identified based on their significantly smaller body size. We estimated the perpendicular distance of the frog to the transect using a distometer (Leica Geosystems D1 laser distance meter; range: 0.2 - 40 m; accuracy: ± 2 mm).

**Figure 1.**
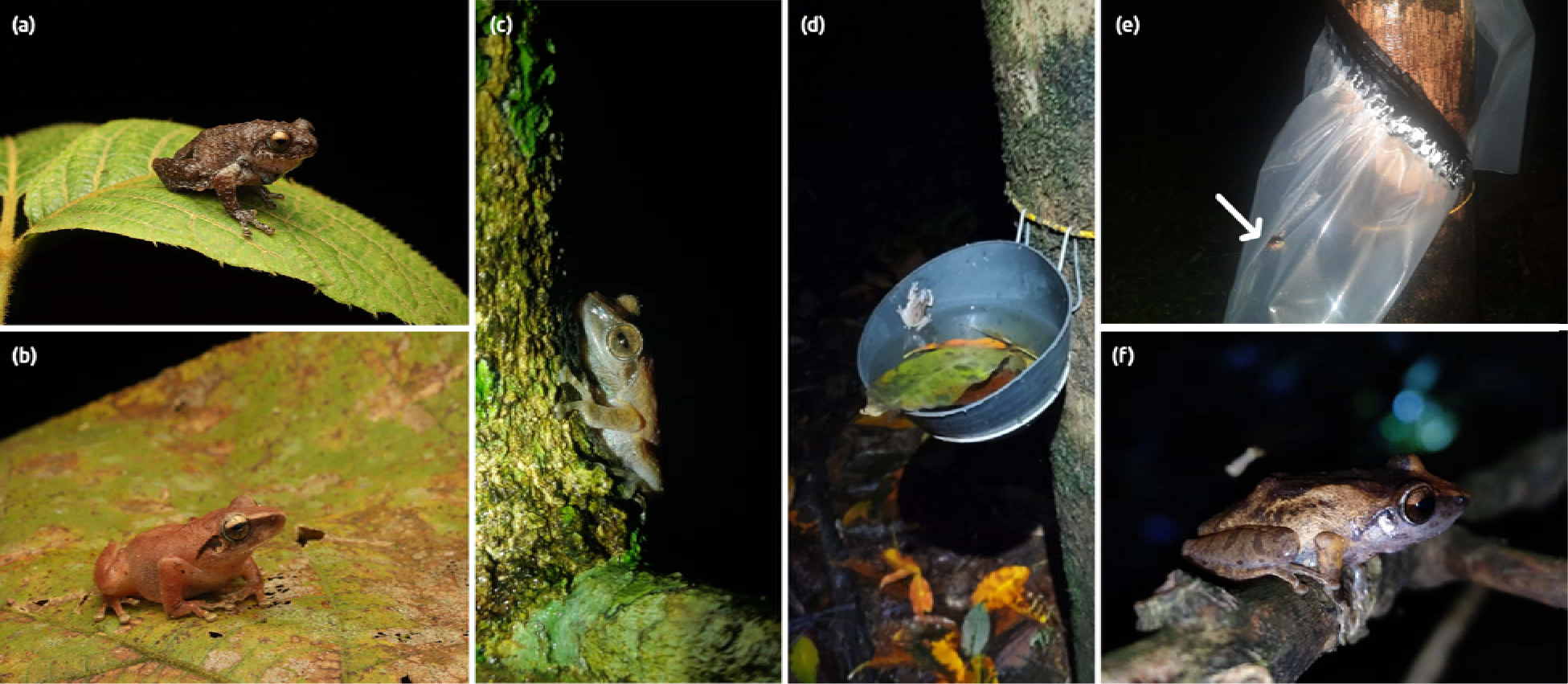
Photographs showing shrub frogs using different substrates, such as (a) *Raorchestes bombayensis* on a leaf in high-elevation forests, (b) *Psuedophilautus amboli* photographed on leaf litter in low-elevation forests, (c) trunk, (d) a plastic bowl used for collecting rubber latex, (e) a plastic sheet used to protect collected latex from rain, and (f) branch. Photographs by Rohit Naniwadekar, Ninad Gosavi and Himanshu Lad.

### Habitat differences across land use types

We measured all the habitat variables in a 5m × 5m plot at every 50 m interval, i.e. six plots per transect across land use categories. We measured the leaf litter depth using a metal ruler at four corners of a 1m^2^ plot. We counted the number of shrubs (woody stem, girth at breast height (GBH) < 15cm) and number of trees (GBH > 15cm) in each 25m^2^ plot. Additionally, we visually estimated the percent cover of bare ground, grass, undergrowth, and shrubs in the 25m^2^ plot.

### Microhabitat use by shrub frogs

We recorded perch substrate and height for every shrub frog sighting during and/or outside transect sampling. We classified the perch substrate into trunk, branch, leaf, fallen branch, leaf litter and plastic. Plastic refers to frogs observed in rubber plantations perched on plastic bowls used for collecting rubber latex and on the plastic sheets used to protect collected latex from rain (Fig SX). We recorded perch height using the Leica distometer.

### Body size of *Pseudophilautus amboli* across land use types

We measured the snout-vent length (SVL) of calling males of *Pseudophilautus amboli* across the land use categories (Cashew = 13 individuals, Rubber =16 individuals, low-elevation Forest = 15 individuals) using a Mitutoyo 500-196-20 Digital Vernier Caliper (precision = 0.01mm). Previous study has demonstrated that measurements of 14-17 individuals per group are sufficient to detect intra-annual variation in body condition for *Leptodactylus latrans* (Brodeur et al., 2020). After taking the measurements in the field, we immediately released the frogs back at the same location where they were collected. Within each category, we measured the SVL of frogs from different transects spread across the study area to ensure that no biases were induced by spatial variation in body lengths. All frogs were measured within a 15-day period in August to ensure that body size was not influenced by temporal variation.

## ANALYSIS

### Habitat differences across land use types

To understand how the microhabitat availability varied across the four land-use categories, we performed unconstrained principal coordinate analysis (PCoA) using the Bray-Curtis dissimilarity index using *cmdscale* function of default package ‘stats’ and vegdist, and *envfit* functions of the package ‘vegan’ (Oksanen et al., 2022). We visualised the differences in microhabitat data across the land use categories (low- and high-elevation forests, cashew and rubber) in a biplot along with environment vectors representing the microhabitat features onto the ordination. We then used the ‘betadisper’ function to assess the multivariate homogeneity of group dispersions.

### Density estimation

We used the ‘Distance’ package in R to estimate the densities of adult male and juvenile *P. amboli* and *R. bombayensis* (Miller et al., 2019). Information on sampling effort (pooled across temporal replicates for a transect), perpendicular distance, and cluster size were used to estimate individual densities of shrub frogs. Given our expectation of differing detection probabilities between species and life stages (adults and juveniles), we analysed the data separately for the two species of shrub frogs and adult males and juveniles (Table S2). Considering potential variations in detection functions across land use types, we implemented the multi-covariate distance sampling (MCDS) protocol with land use category as a covariate to estimate densities across land use types using the functions ‘ds’ and ‘dht2’ as implemented by package ‘Distance’ in R (Miller et al., 2019). We employed both the half-normal and hazard rate models to estimate detection probability, and the model with the lowest Akaike’s Information Criteria (AIC) value was considered the best.

### Microhabitat use by shrub frogs

We employed the chi-square test of independence to determine differences in the relative proportions of detections for 1) *P. amboli* across three land use types and 2) between *P. amboli* and *R. bombayensis* in forest habitats. Since perch height data were not normally distributed (Shapiro-Wilk Test; *W* = 0.919, *p* < 0.001), we performed the non-parametric Kruskal-Wallis test to compare the perch height of calling males across land use categories. For post-hoc pairwise differences between categories, we used Dunn’s test using the function ‘dunnTest’ as implemented in package ‘FSA’ in R (Ogle et al., 2024).

### Body size of *Psuedophilautus amboli* across land use types

Snout-vent length (SVL) data of male *P. amboli* was normally distributed (Shapiro-Wilk Test; *W =* 0.988, *p* = 0.92). Thus, we used a general linear model (with a Gaussian error distribution) to determine differences in body size among male frogs across the three land use types. We used Tukey’s post-hoc test to determine pairwise differences in SVL across land use types.

## RESULTS

### Habitat differences across land use types

The first two axes of PCoA explained 27.4% (PCo1: 17.8%; PCo2: 9.6%) of the variation in habitat data, indicating differences in habitat structure across land use categories (Fig. 2). Forest plots exhibited higher tree and shrub density, ground cover, and litter depth, contrasting with cashew and rubber plots that tended to have greater grass cover.

**Figure 2.**
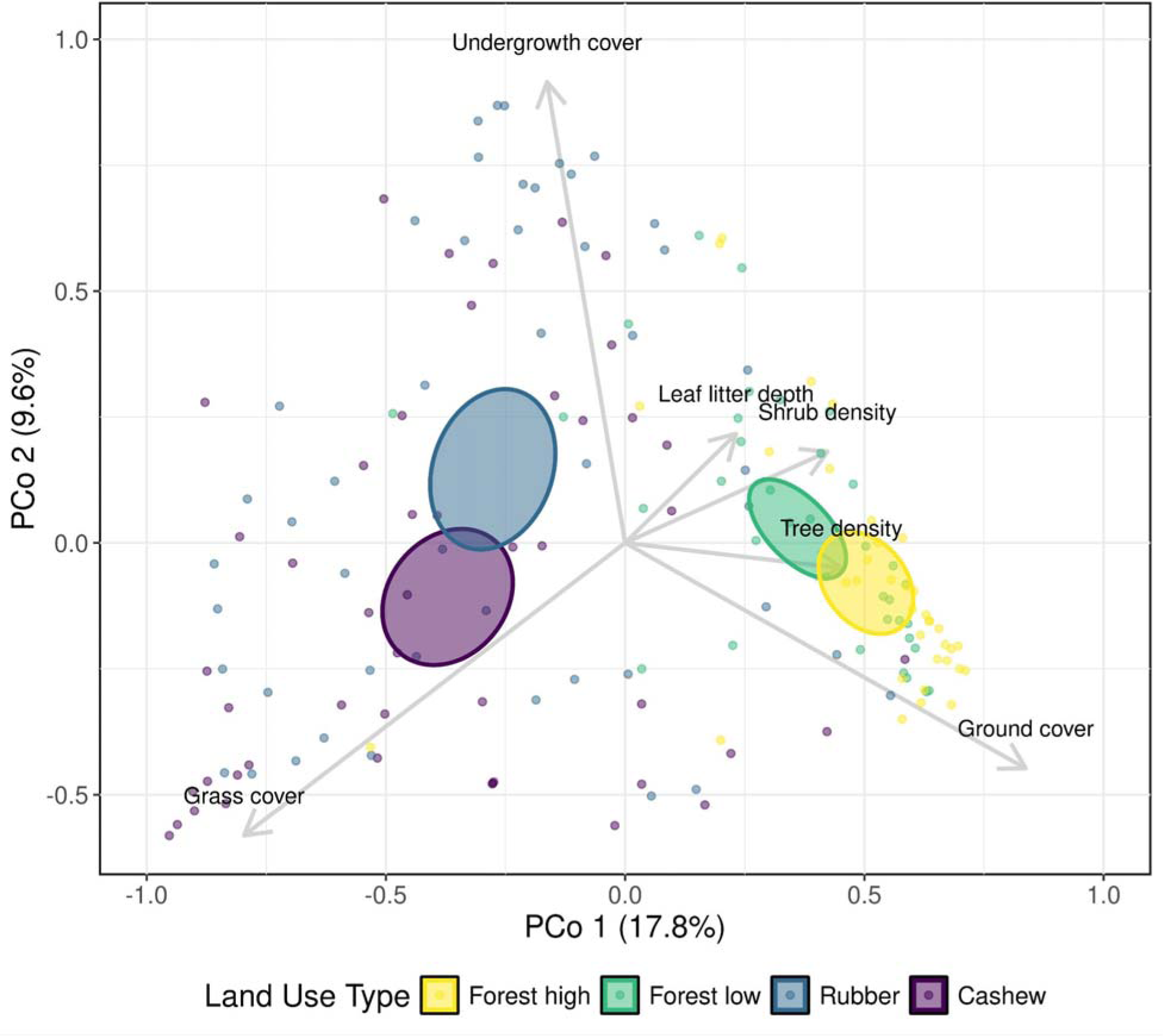
Principal Coordinate Analysis ordination plot showing different habitat variables based on the Bray-Curtis distance measure. The percentages in parentheses on the x- and y-axes indicate the variation explained by the respective axes. The 95% confidence-level ellipsoids representing the four land use types highlight differences in habitat structure among them.

### Densities of shrub frogs

We had 523 detections of *P. amboli* and 167 detections of *R. bombayensis* (Table S2). In low elevations, we had at least 245 detections each of adult males and juveniles of *P. amboli*, while in high elevations, we had at least 67 detections each of adult males and juveniles of *R. bombayensis* (Table S2). However, we had very few detections of *P. amboli* in high-elevation sites (8 individuals) and *R. bombayensis* in low-elevation sites (11 individuals). Importantly, we had very few detections of adult female shrub frogs (*P. amboli*: 9 individuals; *R. bombayensis*: 2 individuals). Consequently, we did not estimate densities of *P. amboli* for high elevations, *R. bombayensis* for low elevations, and females across elevations and land use categories. Notably, the adult female to adult male ratios were extremely low for both *P. amboli* (0.03) and *R. bombayensis* (0.03), indicating highly skewed, male-biased sex ratios in these species.

A summary of the different models we implemented for estimating shrub frog densities is provided in Table S3. While the hazard rate key function best fitted the data for the adult *P. amboli* and juvenile *R. bombayensis*, the half-normal key function best fitted the data for juvenile *P. amboli* and adult *R. bombayensis* (Table S3). In all cases, the models fit the data well (*p* > 0.05) (Table S3). For *P. amboli,* a model with detection probability modeled as a function of land-use type had the least AIC value (Table S3), justifying the use of distance sampling and MCDS for density estimation.

The mean densities of adult male and juvenile *P. amboli* were notably lower in cashew plantations compared to forests and rubber plantations, as inferred by 95% CI not overlapping the means (Fig. 3; Table S4). Specifically, the mean densities were 2.4 times higher in the forests and 3.7 times higher in the rubber plantations than in cashew plantations (Fig. 3). Juvenile densities were 15.1 times higher in the forests and 17 times higher in the rubber plantations than the cashew plantations (Fig. 3). Furthermore, the ratio of juveniles to adults was 2.4 in forests and 1.7 in rubber, whereas it was only 0.4 in cashew plantations, suggesting poor recruitment of *P. amboli* in cashew plantations.

**Figure 3.**
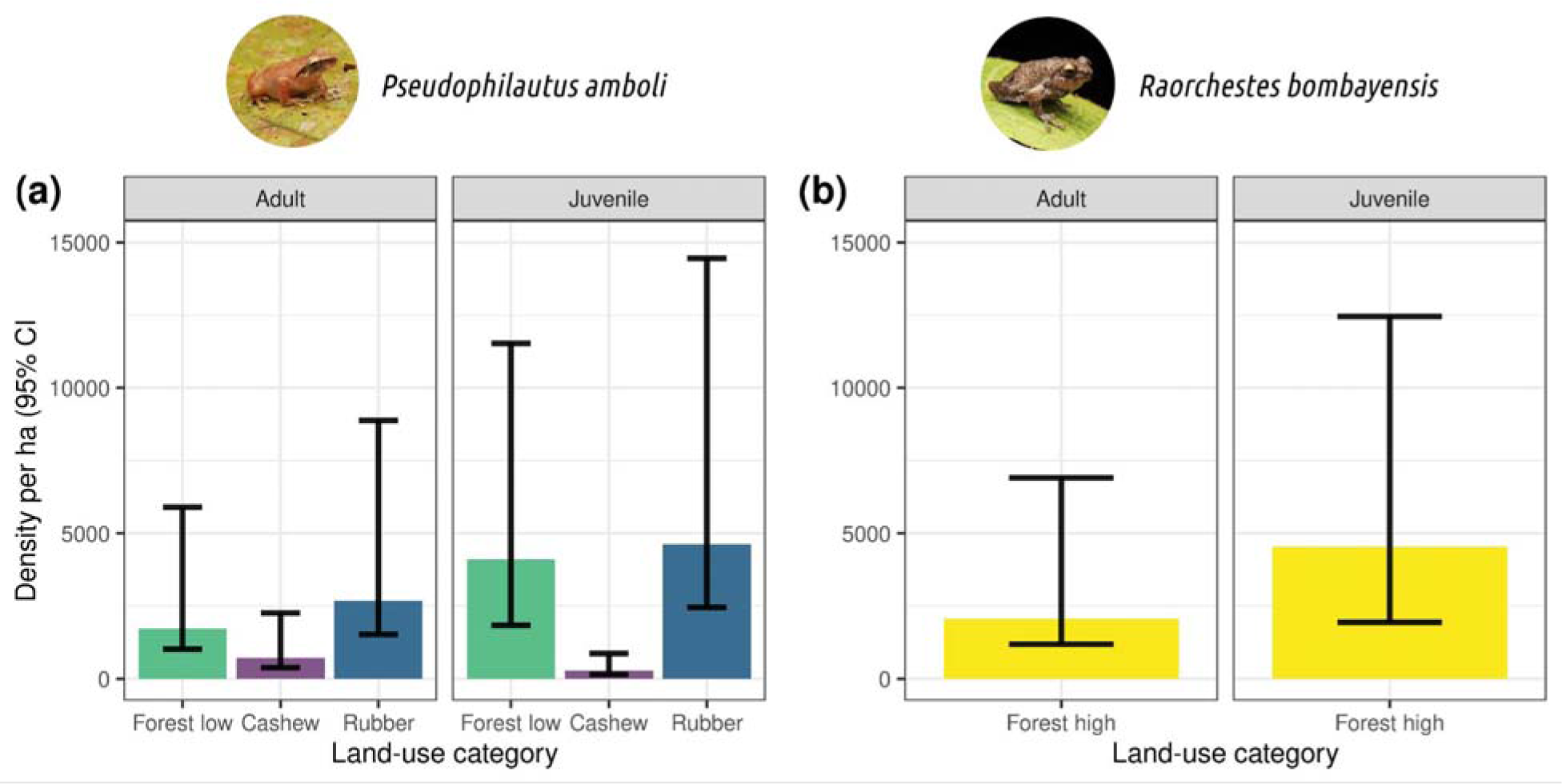
Individual densities (per ha) (± 95% CI) of the two species of shrub frogs with the values of the mean indicated on top of the error bars. Due to limited detections, we did not estimate the *R. bombayensis* in low elevations and *P. amboli* in high elevations.

### Microhabitat-use across land-use types

There were significant differences in the relative use of different substrates by *P. amboli* across the land-use types (□*^2^*= 87.373, df = 10, *p* < 0.001). Notably, we had very few detections of shrub frogs on the main trunk in cashew and rubber plantations (Fig. 4A). In rubber plantations specifically, we had at least twice as many detections of Amboli shrub frogs on plastic sheets than on the main trunk (Fig. 4A). Additionally, there were significant differences in the relative use of different substrates by *P. amboli* and *R. bombayensis* in forests (□^2^= 10.851, df = 4, *p* = 0.029). Most of our *R. bombayensis* detections were on leaves, with relatively fewer detections on branches and trunks (Fig. 4B).

**Figure 4.**
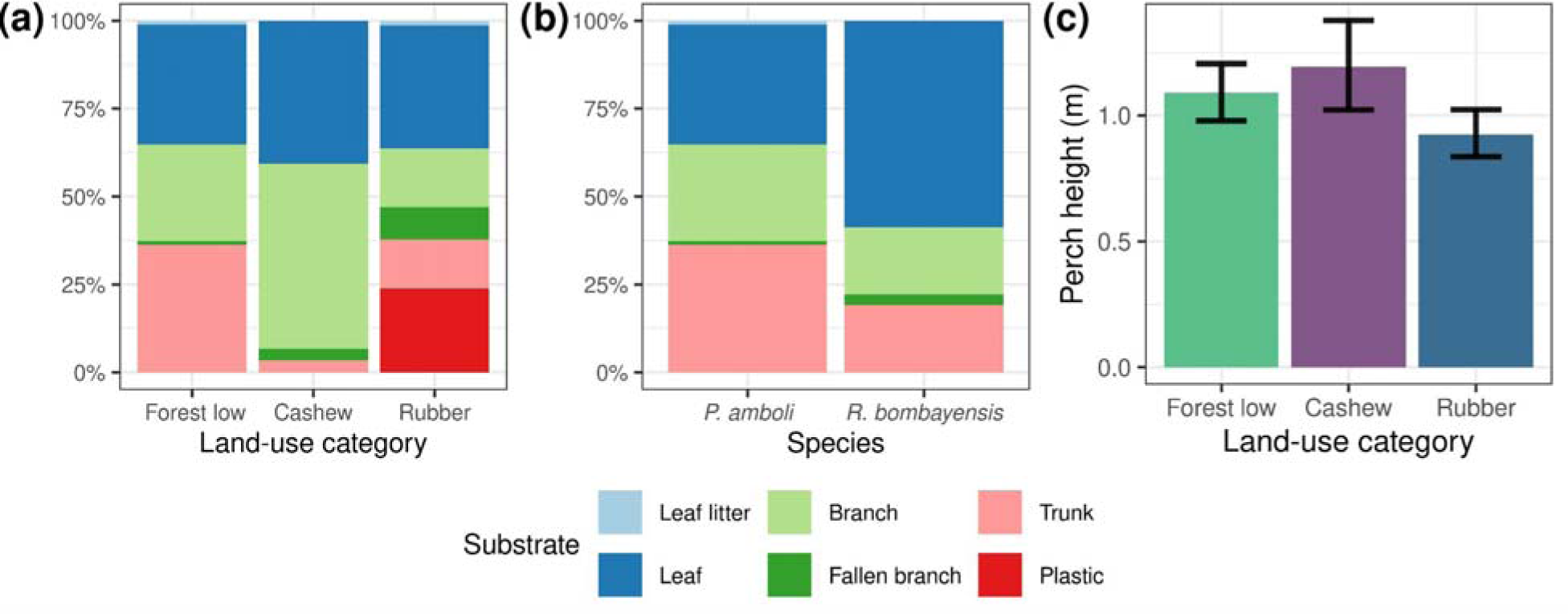
**(a)** A comparison of substrates used by *Pseudophilautus amboli* in cashew, rubber and low-elevation forests, (b) a comparison of substrates used by *P. amboli* in low-elevation forests and *R. bombayensis* in the high-elevation forests; (c) mean (bootstrapped 95% CI) values of perch heights of *Pseudophilautus amboli* male frogs (n=256) across land use categories. The perch height used by frogs in rubber was significantly lower than in low-elevation forests and cashew.

The perch height of *P*. *amboli* male frogs varied significantly across land-use types (*χ^2^* = 14.952, df = 2, *p* < 0.001). The perch height (mean ± SE) in the forest (1.12 ± 0.05) was 1.2 times higher, and in cashew (1.18 ± 0.08) was 1.3 times higher than in rubber (0.92 ± 0.04). Dunn’s test indicated a significant difference in perch height between low-elevation forests and rubber (*p* = 0.002) and cashew and rubber (*p =* 0.01). (Fig. 4c; Table S5)

### Snout-vent length of frogs across land-use types

The mean (± SE) snout-vent length of frogs across the three land-use categories were as follows: Cashew: 27.3 mm (± 0.5), Rubber: 26.5 mm (± 0.3), Forest (low): 27.8 mm (± 0.3). The snout-vent length of male frogs was significantly smaller in rubber plantations than in low-elevation forests (*F_2,41_* = 3.373, *p* = 0.044) (Fig. 5, Table S6 and S7).

**Figure 5.**
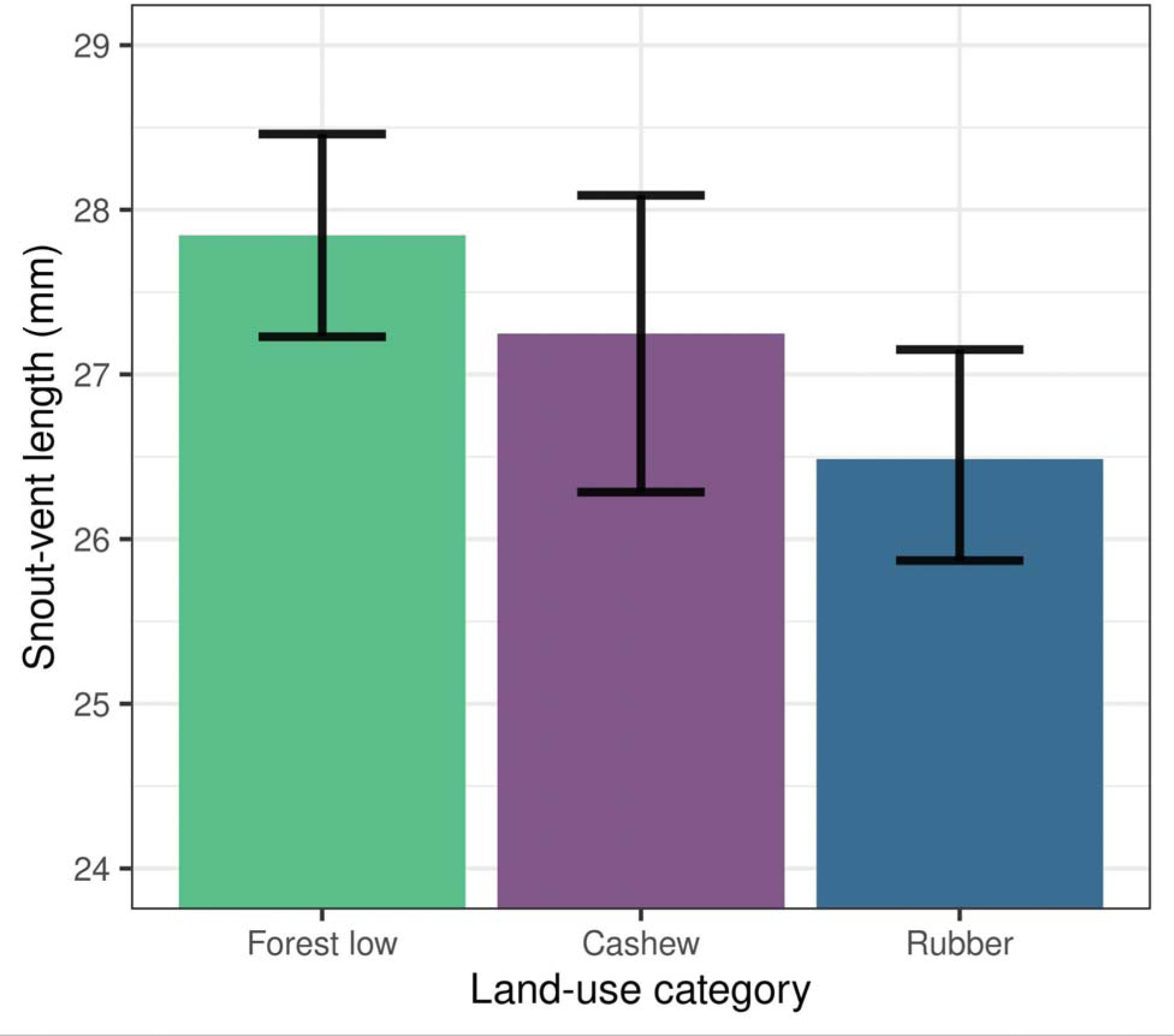
Mean (bootstrapped 95% CI) values of snout-vent length of adult males of *Pseudophilautus amboli* (n = 44) across land use categories. The snout-vent length of frogs in rubber plantations was lower than those in low-elevation forests.

## DISCUSSION

We compared densities and microhabitat use across low and high elevation sites and across different land use types for two endemic shrub frog species, and found differences in elevation, population demography, microhabitat use and body condition of shrub frogs across land use types. We found that *Pseudophilautus* prefers lower elevations, while *Raorchestes* is more abundant at higher elevations in this region. Moreover, these two species differ in substrate preferences. Notably, the responses of *Pseudophilautus* to different forms of agroforestry varied. Cashew had lowered densities, poor recruitment, and altered microhabitat usage, while rubber had altered microhabitat use without impacting *Pseudophilautus* densities. Furthermore, adult male frogs in rubber were significantly smaller than those in forests, indicating potential implications on body condition of frogs. Information on population assessments of shrub frogs has been identified as an important knowledge gap (IUCN SSC Amphibian Specialist Group, 2021), and this study attempts to fill that gap by generating among the first estimates of shrub frog densities in the Western Ghats. We demonstrate that different forms of agroforestry may differ vastly in their impacts on endemic species, emphasising the importance of separately evaluating the effects of each agroforestry type. This study also highlights extremely skewed, male-biased sex ratios of shrub frogs, an aspect that needs careful consideration in species assessments.

### Altered microhabitat use in modified habitats

Interestingly, we found differences in microhabitat use of *Pseudophilautus* in different agroforestry ecosystems. In forests, *Pseudophilautus* were detected on trunks, branches and on leaves; however, in cashew plantations we barely detected them on the main trunk since cashew trees rarely have straight trunks. Interestingly, in rubber plantations, *Pseudophilautus* were observed using novel microhabitats, as they were more commonly found on plastic sheets covering the cups set up to collect the rubber latex than on the trunk. This indicates the ability of the shrub frogs to adapt to modified conditions. The plastic sheets may protect them from excessive rain and predators like arboreal snakes. Moreover, the plastic cover can also influence the acoustic signals. These aspects require further exploration.

### Impact of cashew and rubber plantations

Amphibian abundances are lower in agroforestry ecosystems (Cervantes-López & Morante-Filho, 2024). However, we found that densities of shrub frogs were specifically lower in cashew plantations, with no significant decrease observed in rubber plantations. In contrast to a previous study from the landscape that reported *Pseudophilautus* as an indicator species of cashew plantations (Komanduri, Sreedharan, & Vasudevan, 2023), we found that cashew habitats acted as sink habitats for the species. However, looking at the poor recruitment of the species, *Pseudophilautus* is unlikely to be an indicator of cashew. Cashew plantations have been reported to have significantly lower relative abundance of *P. wynaadensis* in southern Western Ghats (Syamili & Nameer, 2018).

Comparisons of population demography of frogs across land use types is lacking, particularly from south Asian wet tropics, a hotspot for amphibians. In our case, not only were the densities of *Pseudophilautus* significantly lower in cashew plantations, but the juvenile to adult ratios were less than one, indicating poor recruitment in cashew plantations. This was in contrast to forests and rubber plantations, where the juvenile-to-adult ratios were more than one. Most cashew plantations in the landscape are relatively small in size and are surrounded by secondary forests (Rege, Warnekar, & Lee, 2022). These forests could act as a source for adults. The reasons for poor recruitment of shrub frogs in cashew plantations need to be determined in future studies. It is critical to ascertain at which stage filtering occurs; for instance, whether egg laying is impacted or if the mortality of eggs and juveniles is higher in cashew plantations. Generally, the decrease in favourable habitats in agroforestry plantations are speculated to be a driver for reduced abudances of frogs (Cervantes-López & Morante-Filho, 2024). However, both rubber and cashew plantations had lower tree and shrub density, as well as reduced leaf litter depth. Therefore, these factors are unlikely to be the driving factors behind the observed decline in cashew plantations at least in our case. Cashew plantations had greater grass cover and lower understorey cover. Whether these factors influence availability of egg laying sites for shrub frogs needs to be determined in future studies.

Pesticides are known to influence amphibian richness (Rathod & Rathod, 2013; Wanger et al., 2023). Differences in unmeasured variables, like herbicides and pesticides use, can also explain differences in shrub frog abundance across land use categories. Herbicides are extensively used in cashew plantations and not as much in rubber plantations based on conversations with local researchers and cashew farmers. While herbicides and pesticides are not applied during the monsoon period when the frogs breed (personal observations; Vishal Sadekar personal communication), it is likely that remnants in the environment may influence frog abundance in cashew plantations, an aspect that requires further investigation. Moreover, the undergrowth in cashew is periodically cleared during the monsoon. Future studies need to disentangle the relative role of herbicides, pesticides, and undergrowth removal using experimental approaches.

Rubber plantations are known to support high amphibian diversity, likely due to heterogeneous and stable tree structure and high leaf litter availability (Paoletti et al., 2018). In our case, rubber plantations had lower shrub and tree density and leaf litter compared to forests, yet they had similar densities of adults and juveniles to forests. Rubber plantations in central Western Ghats were found to be dominated by a few species that are relatively ubiquitous, including *Pseudophilautus* and *R. tuberohumerus* (Sankararaman et al., 2021). The relative abundance of *P. wynaadensis*, one of the most common amphibians detected in the community, was higher in rubber plantations than in other agroforestry land uses (Syamili & Nameer, 2018). However, both these studies lacked data from reference ecosystems. Despite similar densities of *Pseudophilautus* in rubber and forests, its body size was significantly smaller than forests. Previous studies have shown reduced body size in leaf-litter dwelling *Rhinella ornata* in fragmented habitats (Steinicke et al., 2015), and *Phyllomedusa tarsius* in disturbed habitats (Neckel-Oliveira & Gascon, 2006). *Eleutherodactylus antillensis* and *E. coqui*, two directly developing species similar to our case, have been found to show reduced body size in disturbed habitats (Delgado-Acevedo & Restrepo, 2008). Decrease in body size can result from inadequate resources, negative effects of contamination, or reduced longevity with consequences for reproductive output and success (de L. Bionda et al., 2018; Jennette, Snodgrass, & Forester, 2019; Cogălniceanu et al., 2021). Future research is required to determine drivers and consequences of reduced body size for shrub frog populations, including survival probabilities and clutch size.

### Implications of male-biased sex ratios

While Rhacophorids have been reported to have even and male-biased sex ratios, we found that the sex ratios were extremely male-biased in shrub frogs (3 females for every 100 males). Given that we detected significant number of juveniles (Table S2), which do not call and are much smaller than males, we are confident that these biased sex ratios are not an outcome of detection bias. We argue that the high densities of shrub frogs are maintained by a small number of females (around 50 females/ha to around 1700 males/ha of *Psuedophilautus*). If the populations of females, which occur in much lower densities than males, are impacted, it will result in the shrub frog populations crashing. Moreover, we demonstrate that *Pseudophilautus*, which has been thought to be a generalist and resilient species (Sankararaman et al., 2021), is negatively impacted by cashew plantations, which are rapidly increasing in the landscape (Rege & Lee, 2022) and currently occupy almost a third of the geographic area of the Dodamarg and Sawantwadi Taluks where the study was conducted. This is an important consideration for the IUCN Redlist Status assessment, which recently downlisted these two endemic species from Threatened category to Least Concern (IUCN SSC Amphibian Specialist Group, 2021). We urge that given this novel information on extremely skewed sex ratios, the vulnerability of *Pseudophilautus* to cashew plantations and other impacts on microhabitat use and body size, differences in densities across elevational range, patchy distribution even in forest habitats (Table S8), and the relatively narrow geographic ranges of the species, the downlisting of these species to Least Concern should be reconsidered and they should be at least classified as “Near Threatened”. This is something that even the Geospatial Conservation Assessment Tool (https://geocat.iucnredlist.org/) for the two species recommends just based on assessing the extent of occurrence information of the location data from Global Biodiversity Information Facility (GBIF).

Moreover, there is also a need to determine the drivers of the skewed sex ratios and its consequences for the shrub frog populations. Skewed sex ratios can be an outcome of sex-biased mortality or sex reversal (Lambert, Ezaz, & Skelly, 2021). Sex reversal, especially from females to males, has been shown to increase with increasing anthropogenic disturbances, including habitat fragmentation, degradation and rising temperatures (Nemesházi et al., 2020).

### Distance sampling for shrub frog population estimation

The need for increased population monitoring has been identified as a crucial measure for threatened amphibians (Luedtke et al., 2023; Re:wild, Synchronicity Earth, 2023). A method that is non-invasive, replicable, easy to implement in the field, factors for variation in detection probability across habitats, and reliably estimates population densities is vital. With more than 60 species of shrub frogs in the Western Ghats, over 50% of which are threatened (Re:wild, Synchronicity Earth, 2023), monitoring their populations across land use categories is critical. With modest effort, we obtained the necessary detections not only of adult male shrub frogs but also of juveniles, that enabled us to estimate shrub frog densities reliably. The significant number of detections of juveniles, which are smaller and do not call, allows studying population demography as well. Moreover, distance sampling is a non-invasive method that allows monitoring of shrub frogs without handling the animals. The wide confidence intervals and high coefficient of variation are an outcome of patchy distribution of shrub frogs even within each land use (Table S8). Thus, finer stratification within a habitat may help reduce coefficient of variation and increase precision in density estimates.

## Conclusions

Our study highlights significant impacts of agroforestry plantations on the population demography, habitat use and body condition of threatened frogs. We demonstrate distinct responses of shrub frogs to different forms of agroforestry plantations, highlighting the importance of utilising different measures to determine species responses to different forms of agriculture (Jithin, Rane, Watve, & Naniwadekar, 2023). Considering the elevational differences in species abundances, highly skewed sex ratios, the vulnerability of shrub frogs to expanding cashew plantations, we caution against the drastic downlisting of endemic and narrow-range species, particularly in the absence of critical ecological information that can inform such decisions. For the two focal species recently downlisted to Least Concern, we advocate for reconsideration to at least Near Threatened status in light of the findings presented here. Future research should focus on determining the drivers of skewed sex ratios that may be influenced by chemicals used in agriculture and increasing temperatures (Nemesházi et al., 2020; Lambert, Ezaz, & Skelly, 2021), and their consequences for shrub frog populations in this changing landscape.

## ACKNOWLEDGEMENTS

This study was funded by the On the Edge Conservation Grant (UK). We thank Maharashtra Forest Department, especially Sunil Limaye (CWLW), for giving us the necessary permits (Letter No. Desk-22(8)/WL/Research/CR-53(20-21) /3361/22-22). We thank Vishal Sadekar, Siddharth Biniwale, Praveen Desai, Parag Rangnekar, Gajanan Shetye, Vijay Karthick, Sudhakar Desai, Rohan Thakur, Hemant Ogale, and Kaka Bhise for their immense support during fieldwork. We thank Anand Osuri and Kulbhushansingh Suryawanshi for their valuable feedback and support.

## AUTHORSHIP STATEMENT

RN conceived the ideas and designed the methodology with inputs from HL and NG; HL and NG collected the data; HL and RN analysed the data with inputs from VJ; RN, HL, and VJ led the writing of the manuscript with inputs from NG. All authors contributed critically to the drafts and gave final approval for publication.

## OPEN RESEARCH STATEMENT

All codes and statistical packages used in this study are cited in the manuscript and publicly available. The adapted codes used for the analyses and the source data will be uploaded on Zenodo on acceptance.

