## Supplementary material for "Effects of land use change and elevation on endemic shrub frogs in a biodiversity hotspot": Table S

*This file includes:*

**Table S1.** Summary of sampling effort across the different land-use types.

**Table S2.** Summary of detections of the two shrub frog species across land-use categories

**Table S3.** Summary of models for distance analysis to estimate densities of adults and juveniles

**Table S4.** Densities of adult and juvenile *Pseudophilautus amboli* and *Raorchestes bombayensis* across different elevation zones and land use types

**Table S5.** Summary of Dunn’s post-hoc tests determining pairwise differences in perch height of calling males of *P. amboli* across the different land-use types.

**Table S6.** Coefficient table as obtained from general linear model analysis that compared snout-vent lengths of adult male *P. amboli* across three land-use types.

**Table S7.** Summary of Tukey’s post-hoc tests determining pair-wise differences in snout-vent length of *P. amboli* across the different land-use types.

**Table S8.** Number of individuals seen during distance sampling across the different land use types.

**Table S1.** Summary of sampling effort across the different land-use types.

| **Land-use type** | **Number of transects** | **Transect length (range) (km)** | **Total effort (km)** |
| --- | --- | --- | --- |
| Cashew | 8 | 0.28-0.30 | 9.22 |
| Rubber | 8 | 0.27-0.30 | 9.39 |
| Forest (low) | 6 | 0.30 | 7.20 |
| Forest (high) | 6 | 0.23-0.30 | 6.48 |
| Total | 28 | 0.23-0.3 | 32.29 |

**Table S2.** Summary of detections of the two shrub frog species across the different land-use categories

| **Species** | **Land-use type** | **Adult female** | **Adult male** | **Juvenile** |
| --- | --- | --- | --- | --- |
| *Pseudophilautus amboli* | Cashew | 3 | 49 | 14 |
|  | Rubber | 5 | 138 | 152 |
|  | Forest (low) | 1 | 74 | 79 |
|  | Forest (high) | 0 | 5 | 3 |
|  | Total | 9 | 266 | 248 |
| *Raorchestes bombayensis* | Cashew | 0 | 1 | 1 |
|  | Rubber | 0 | 3 | 2 |
|  | Forest (low) | 0 | 2 | 1 |
|  | Forest (high) | 2 | 61 | 94 |
|  | Total | 2 | 67 | 98 |

**Table S3.** Summary of models for distance analysis to estimate densities of adults and juveniles of *P. amboli* and *R. bombayensis*. GoF-Goodness of Fit

| **Species** | **Model** | **Covariate** | **AIC** | **ΔAIC** | ***p* (GoF)** | **CV** |
| --- | --- | --- | --- | --- | --- | --- |
| *P. amboli* (adult) | Hazard rate | ~Land-use type | 582.36 | 0 | 0.588 |  |
|  | Half-normal | ~1 | 584.25 | 1.898 | 0.961 |  |
|  | Hazard rate | ~1 | 586.02 | 3.664 | 0.938 |  |
|  | Half-normal | ~Land-use type | Model did not converge | | |  |
| *P. amboli* (juvenile) | Half-normal | ~Land-use type | 565.416 | 0 | 0.200 |  |
|  | Hazard rate | ~Land-use type | 566.606 | 1.189 | 0.516 |  |
|  | Hazard rate | ~1 | 567.117 | 1.701 | 0.945 |  |
|  | Half-normal | ~1 | 569.171 | 3.755 | 0.349 |  |
| *R. bombayensis* (adult) | Half-normal | ~1 | 125.994 | 0 | 0.207 |  |
|  | Hazard rate | ~1 | 126.407 | 0.413 | 0.233 |  |
| *R. bombayensis* (juvenile) | Hazard rate | ~1 | 168.353 | 0 | 0.856 |  |
|  | Half-normal | ~1 | 169.278 | 0.924 | 0.574 |  |

**Table S4.** Densities of adult and juvenile *Pseudophilautus amboli* and *Raorchestes bombayensis* across different elevation zones and land use types and associated standard, coefficient of variation (CV) and confidence intervals (CI)

| **Species** | **Land-use type** | **Density** | **SE** | **CV (%)** | **95% LCI** | **95% UCI** |
| --- | --- | --- | --- | --- | --- | --- |
| *Pseudophilautus* Adult | Forest low | 1716 | 649 | 37.8 | 704 | 4180 |
|  | Cashew | 723 | 240 | 33.1 | 341 | 1533 |
|  | Rubber | 2676 | 1043 | 39.0 | 1154 | 6205 |
| *Pseudophilautus* Juvenile | Forest low | 4108 | 1027 | 25.0 | 2274 | 7421 |
|  | Cashew | 273 | 108 | 39.7 | 125 | 596 |
|  | Rubber | 4631 | 1534 | 33.1 | 2183 | 9823 |
| *Raorchestes* Adult | Forest high | 2056 | 743 | 36.1 | 871 | 4854 |
| *Raorchestes* Juvenile | Forest high | 4531 | 1084 | 24.0 | 2592 | 7920 |

**Table S5.** Summary of Dunn’s post-hoc tests determining pairwise differences in perch height of calling males of *P. amboli* across the different land-use types.

|  | ***Z*** | ***p*-adjusted** |
| --- | --- | --- |
| Forest (low)-Cashew | 0.11244 | 1 |
| Forest (low)-Rubber | 3.38788 | 0.00211 |
| Cashew-Rubber | 2.88576 | 0.01171 |

**Table S6.** Coefficient table as obtained from general linear model analysis that compared snout-vent lengths of adult male *P. amboli* across three land-use types.

|  | **Estimate** | **SE** | ***t*-value** | ***p*** |
| --- | --- | --- | --- | --- |
| Intercept: Forest (low) | 27.844 | 0.364 | 76.561 | <0.001 |
| Cashew | -0.596 | 0.543 | -1.097 | 0.279 |
| Rubber | -1.356 | 0.523 | -2.594 | 0.013 |

**Table S7.** Summary of Tukey’s post-hoc tests determining pair-wise differences in snout-vent length of *P. amboli* across the different land-use types.

|  | **Difference** | **Lower 95% CI** | **Upper 95% CI** | ***p*-adjusted** |
| --- | --- | --- | --- | --- |
| Cashew-Forest (low) | -0.596 | -1.917 | 0.725 | 0.521 |
| Rubber-Forest (low) | -1.356 | -2.628 | -0.085 | 0.034 |
| Rubber-Cashew | -0.761 | -2.101 | 0.58 | 0.361 |

**Table S8.** Number of individuals seen during distance sampling across the different land use types.

| **Land use type** | **Transect id** | **Adult female** | **Adult male** | **Juvenile** | **Sampling effort** |
| --- | --- | --- | --- | --- | --- |
| Forest (low) | FL1 | 1 | 32 | 17 | 1.2 |
| Forest (low) | FL2 | 0 | 8 | 17 | 1.2 |
| Forest (low) | FL3 | 0 | 19 | 7 | 1.2 |
| Forest (low) | FL4 | 0 | 6 | 5 | 1.2 |
| Forest (low) | FL5 | 0 | 2 | 24 | 1.2 |
| Forest (low) | FL6 | 0 | 7 | 9 | 1.2 |
| Cashew | CT1 | 0 | 3 | 1 | 1.2 |
| Cashew | CT2 | 0 | 1 | 1 | 0.9 |
| Cashew | CT3 | 0 | 1 | 0 | 1.2 |
| Cashew | CT4 | 0 | 2 | 3 | 1.2 |
| Cashew | CT5 | 0 | 12 | 2 | 1.2 |
| Cashew | CT6 | 1 | 5 | 0 | 1.12 |
| Cashew | CT7 | 1 | 16 | 2 | 1.2 |
| Cashew | CT8 | 1 | 9 | 5 | 1.2 |
| Rubber | RT1 | 2 | 41 | 40 | 1.08 |
| Rubber | RT2 | 0 | 44 | 50 | 1.2 |
| Rubber | RT3 | 0 | 7 | 0 | 1.2 |
| Rubber | RT4 | 1 | 8 | 7 | 1.2 |
| Rubber | RT5 | 1 | 9 | 23 | 1.2 |
| Rubber | RT6 | 0 | 7 | 9 | 1.17 |
| Rubber | RT7 | 0 | 17 | 16 | 1.2 |
| Rubber | RT8 | 1 | 5 | 7 | 1.14 |
